# Dissecting the genetic architecture of shoot growth in carrot (*Daucus carota* L.) using a diallel mating design

**DOI:** 10.1101/115519

**Authors:** Sarah D. Turner, Paul L. Maurizio, William Valdar, Brian S. Yandell, Philipp W. Simon

**Affiliations:** Department of Horticulture, University of Wisconsin-Madison, 1575 Linden Drive, Madison, WI 53706, USA; Curriculum in Bioinformatics and Computational Biology, University of North Carolina, Chapel Hill, NC 27599, USA; Department of Genetics, University of North Carolina, Chapel Hill, NC 27599; Department of Statistics, University of Wisconsin-Madison, 1300 University Avenue, Madison, WI 53706, USA; USDA-ARS Vegetable Crops Research Unit, 1575 Linden Drive, Madison, WI 53706, USA

**Keywords:** Diallel, Bayesian mixed model, genetic architecture, multiple imputation, heterosis

## Abstract

Crop establishment in carrot (*Daucus carota* L.) is limited by slow seedling growth and delayed canopy closure, resulting in high management costs for weed control. Varieties with improved growth habit (i.e. larger canopy and increased shoot biomass) may help mitigate weed control, but the underlying genetics of these traits in carrot is unknown. This project used a diallel mating design coupled with recent Bayesian analytical methods to determine the genetic basis of carrot shoot growth. Six diverse carrot inbred lines with variable shoot size were crossed in WI in 2014. F1 hybrids, reciprocal crosses, and parental selfs were grown in a randomized complete block design (RCBD) with two blocks in CA (2015, 2016) and in WI (2015). Measurements included canopy height, canopy width, shoot biomass, and root biomass. General and specific combining abilities were estimated using Griffing’s Model I. In parallel, additive, inbreeding, epistatic, and maternal effects were estimated from a Bayesian linear mixed model, which is more robust to dealing with missing data, outliers, and theoretical constraints than traditional biometric methods. Both additive and non-additive effects significantly influenced shoot traits, with non-additive effects playing a larger role early in the growing season, when weed control is most critical. Results suggest that early season canopy growth and root size express hybrid vigor and can be improved through reciprocal recurrent selection.

**Article Summary:** Breeding for improved competitive ability is a priority in carrot, which suffers yield losses under weed pressure. However, improvement and in-depth genetic studies for these traits relies on knowledge of the underlying genetic architecture. This study estimated heritable and non-heritable components of carrot shoot growth from a diallel mating design using a Bayesian mixed model. Results directly contribute to improvement efforts by providing estimates of combining ability, identifying a useful tester line, and characterizing the genetic and non-genetic influences on traits for improved competitive ability in carrot.

## INTRODUCTION

Carrots are the 7^th^ most consumed fresh vegetable in the United States, with an annual per capita consumption of 3.9 kg (USDA-ERS 2016) and a production value of $762 million USD in 2015 (USDA-NASS 2016). In the US, the high alpha-and beta-carotene content in carrots is the leading source of dietary provitamin A (Block 1994; Simon et al. 2009), which is essential for healthy immune function, vision, reproduction, and cellular communication (Institute of Medicine, Food and Nutrition Board 2001; Johnson and Russel 2004; Solomons 2012). Despite the economic and dietary importance of carrots, crop establishment and productivity remains limited by erratic germination, slow seedling growth, and delayed canopy closure (Rubatzky et al. 1999). This growth habit, coupled with thin, highly segmented leaf laminae, competes ineffectively with weeds for nutrients, water, and light, resulting in yield losses caused by reductions in root size and marketability (Bellinder et al. 1997; Bell et al. 2000). Furthermore, in a survey of weed competitiveness in 25 crops, carrot had the largest reduction in yield under weed pressure (van Heemst 1985).

To limit yield loss, carrots have an extended critical weed-free period ranging from three to six weeks, during which chemical and hand weeding are necessary (Swanton et al. 2010). Hand weeding, while effective, is disruptive to plant growth and requires intensive labor, with estimated costs exceeding $4000 USD/ha (Bell et al. 2000). For organic systems, which constitute 14.4% of carrot acreage in the US (USDA-ERS 2016), hand weeding is typically the primary method of weed management. Even in conventional systems, few herbicides are labeled for carrots and can only be applied when plants reach a threshold height (e.g. linuron) or have five to six true leaves (e.g. metribuzin), by which point weeds have exceeded control stages (Bellinder et al. 1997).

Cultivars with increased weed competitiveness offer a low cost, nonchemical, and sustainable addition to an integrated weed management program (Pérez de Vida et al. 2006; McDonald and Gill 2009). Improved competitive ability has been linked to traits that increase resource allocation, such as height and biomass accumulation, in other densely planted crops such as maize (Mohammadi 2007; Zystro et al. 2012), rice (Ni et al. 2000; Fischer et al. 2001; Pérez de Vida et al. 2006), wheat (Lemerle et al. 1996; Murphy et al. 2008), and soybean (Jannink et al. 2000; da Silva et al. 2013). While improvement of these traits offers a potential solution for weed management, it is unknown how these traits are inherited in carrot or how they influence marketability (e.g. root biomass accumulation).

The diallel mating design, which consists of pairwise combinations among a group of inbred parents, is a classical breeding approach to identify informative testers, select desirable hybrid combinations, and to determine the primary genetic control for complex traits (Hayman 1954a; Hayman 1954b; Gardner and Eberhart 1966). First introduced to plant breeding by Sprague and Tatum in 1942, the relationships generated in a diallel crossing scheme allow estimation of general (GCA) and specific combining abilities (SCA), which correspond to the proportion of additive and non-additive genetic variation, respectively (Sprague and Tatum 1942; Hayman 1954a; Hayman 1954b; Griffing 1956a; Griffing 1956b). However, the diallel mating design is underutilized in genetic studies due to resource constraints, the complexity of the analysis, and untenable assumptions (e.g. no epistasis; independent distribution of genes among parents) (Baker 1978). These challenges often lead to misinterpretation of the estimates obtained from diallel analysis and have been the subject of much debate in the literature (Baker 1978; Hallauer and Miranda 1981; Christie and Shattuck 1992).

Several methods of diallel analysis have been described, of which the methodology proposed by Griffing (1956a) is one of the most common. This method estimates the significance of GCA, SCA, and reciprocal cross effects using a general linear model approach and can be modified to test interactions of main effects across environments (Lin et al. 1977; Zhang and Kang 1997). However, these traditional methods are not robust in addressing common issues encountered in field experiments, such as missing data, imbalance, and outliers. Missing data for a single cross is often addressed by list-or pair-wise deletion, thereby substantially reducing the number of observations and power of the analysis.

The challenges of missing data and theoretical ambiguity have been addressed by the development of Bayesian methods for diallel analysis, which improve biological interpretability and expand the types of questions that can be addressed (Greenberg et al. 2010; Lenarcic et al. 2012). In this study, we used the methodologies developed by Griffing (1956a) and by Lenarcic et al. (2012) to elucidate the relative importance of genetic parameters for shoot growth in carrots. The primary goals of this work were (1) to estimate the inheritance of shoot growth in carrots as a resource to inform selection strategies, identify useful testers, and assess hybrid performance and (2) to present an applied framework for the analysis of multiple environment data using advanced multiple imputation and Bayesian methods.

## MATERIALS AND METHODS

### Plant material and measurements

Six inbred lines, with canopy heights ranging from short (31.4 cm) to tall (52.8 cm), were selected from the USDA-VCRU carrot breeding program and included L6038, L7550, P0159, Nbh2189, P6139, and B7262 (Table 1, Figure 1, **Figures S1-S2**). Inbred parents were combined in all pairwise combinations for a total of 36 combinations. The resulting F1 progenies, reciprocals, and parental selfs were grown in a randomized complete block design (RCBD) with two blocks. Field sites included the Hancock Agricultural Research Station (Hancock, WI, USA; 2015) and the University of California Imperial County Cooperative Extension Station (Holtville, CA, USA; 2015 & 2016). Carrots were grown on 1.5 meter (m) plots with 1 m between-row spacing.

**Figure 1.**
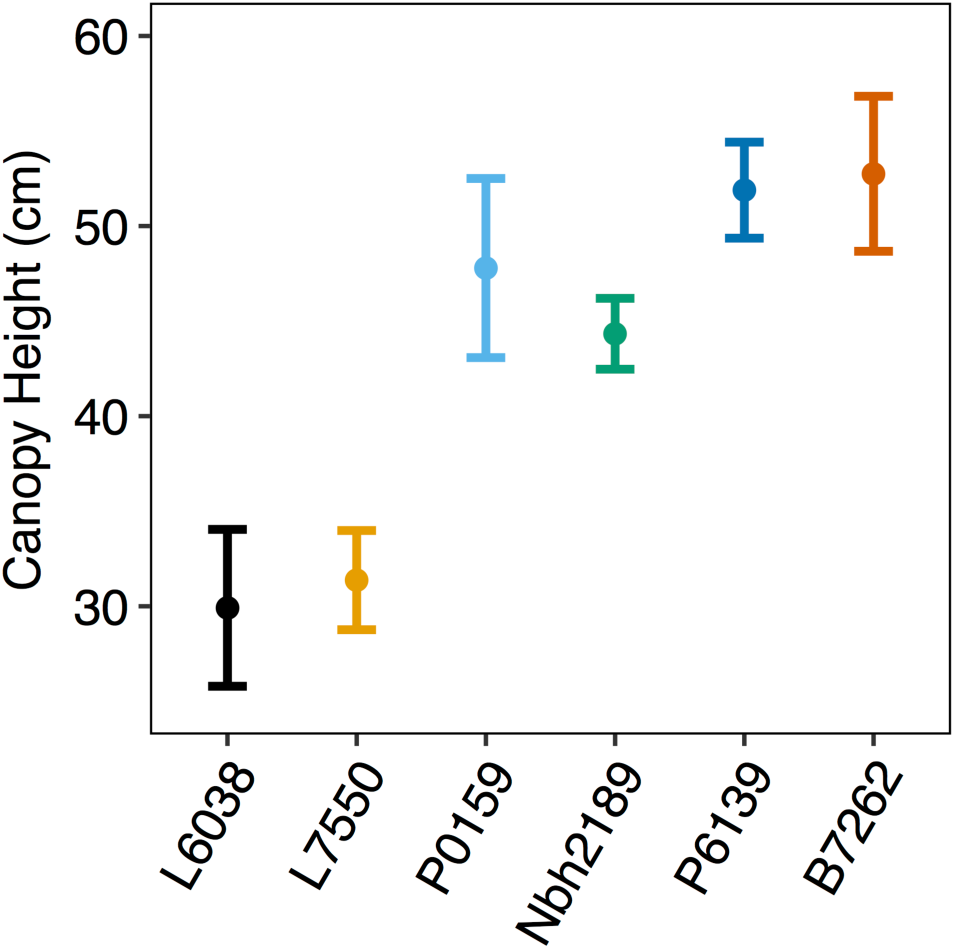
Variation in means and 95% confidence intervals for canopy height (130 DAP) among carrot inbred lines used in this study.

**Table 1.** Carrot traits evaluated, their range among parents, and number of complete observations for each environment in this study.

| Phenotype | Measurement | Unit | Parental range | Data Transformation | Number of complete observations<sup>a</sup> |  |  |
| --- | --- | --- | --- | --- | --- | --- | --- |
|  |  |  |  |  | CA2015 | WI2015 | CA2016 |
| Canopy height | Three points within the plot | Centimeters (cm) | 29.9 – 52.8 | None | 72 | 50 | 69 |
| Canopy width | Three points within the plot | Centimeters (cm) | 41.5 – 61.3 | None | 72 | 50 | 69 |
| Shoot biomass | Fresh and dry | Grams (g) | 6.43 – 21.3 (fresh)<br>1.02 – 3.39 (dry) | ln(x) | 72 | 49 | 68 |
| Root biomass | Fresh and dry | Grams (g) | 29.0 – 64.9 (fresh)<br>4.22 – 8.64 (dry) | ln(x) | 72 | 49 | 68 |
| Shoot to root ratio | Shoot biomass/root biomass (dry) | Grams (g) | 0.23 – 0.64 | ln(x) | 72 | 49 | 68 |

<sup>a</sup> 72 observations possible per environment (36 entries x 2 replications)

Measurements of each trait were taken for three biological replicates per plot and are summarized in Table 1. Canopy height (cm) and width (cm) were measured at midseason, 80 days after planting (DAP), and at harvest, 130 DAP. At harvest, fresh and dry biomass (g) were recorded separately for both shoot and root tissue. For dry biomass, samples were dried at 60°C in a forced-draft oven until reaching a constant weight. A natural log transformation, ln(x), was applied to biomass measurements to make the data distribution symmetric and stabilize the variance.

### Statistical Analyses

Diallel data for each phenotype was analyzed using two complementary approaches: a traditional analysis after Griffing (1956a), which, owing to its requirement that data is complete and balanced, was combined here with a multiple imputation procedure; and the recent Bayesian mixed model decomposition of Lenarcic et al. (2012), performed on the raw (unimputed) data. These are described in detail below. All analyses were performed in R. 3.2.2 (R Core Team 2016) Data and source code are available at https://github.com/mishaploid/carrot-diallel.

### Multiple imputation of missing data for Griffing’s analysis

To compensate for imbalance, missing data (**Figure S3**) was imputed using the Multivariate Imputation by Chained Equations package (R package mice; R/mice) (van Buuren and Groothuis-Oudshoorn 2011), and specifically using that package’s predictive mean matching method (PMM), which is a general purpose, stochastic regression technique that is suitable for numeric data (Little 1988). The predictors used for PMM were chosen based on recommendations in the R/mice documentation, and included male parent, female parent, cross, year, replication, planting density, and numeric measurements with complete data. The values imputed by the PMM were generated by running its associated Markov chain Monte Carlo (MCMC) sampler until it reached a stationary distribution (usually at around 40 iterations; **Figure S4**), and then recording sampled values from a later iteration (e.g. iteration 70). This was repeated *m* = 50 times to generate *m* imputed data sets.

### Griffing’s analysis

Each of the *m* imputed data sets was analyzed using Griffing’s Method I, Model I (1956a), which treats genotypes and blocks as fixed effects and has the base model:

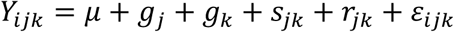

where *μ* is the population mean, *g_j_* and *g_k_* are the GCA effects for the *j*th and *k*th parents, respectively, *s_jk_* is the SCA effect for the cross of the *j*th and *k*th parents (*s_jk_* = *s_kj_*), *r_jk_* is the reciprocal effect for the cross of the *j*th and *k*th parents (*r_jk_* = *–r_kj_*), and *ε_ijk_* is the environmental effect for the *ijk*th observation. Analysis was run using the diallele1 function in the R package plantbreeding (Rosyara 2014), which we modified to include environmental effects and genotype x environment interactions (GxE). The proportion of additive to non-additive genetic variation was estimated from the fixed model by comparing the ratio of mean squares for GCA (
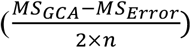
) and SCA (*MS_SCA_ – MS_Error_*), where *n* is the number of parental lines (Baker 1978). Values close to unity suggest higher predictability based solely on GCA. Mean squares and approximate F-tests were pooled following the method proposed by Raghunathan and Dong (2011, unpublished manuscript) (**Table S1**). Estimates for GCA, SCA, and reciprocal effects were combined according to Rubin’s rules (Rubin 1987) and as implemented in R/mice (van Buuren and Groothuis-Oudshoorn 2011).

### Bayesian mixed model for diallel analysis

Raw data for each phenotype (*y_i_*), measured for individuals *i* ∈ {1, …,*n*} with female parent *j* and male parent *k*, were decomposed into additive (*a*), inbreeding (*b*), maternal (*m*), symmetric cross-specific (*v*), and asymmetric cross-specific effects (*w*), as described by Lenarcic et al. (2012) and Crowley et al. (2014) and implemented as a Gibbs sampler in the R package BayesDiallel. Specifically, we used BayesDiallel’s full unsexed model (‘fullu’):

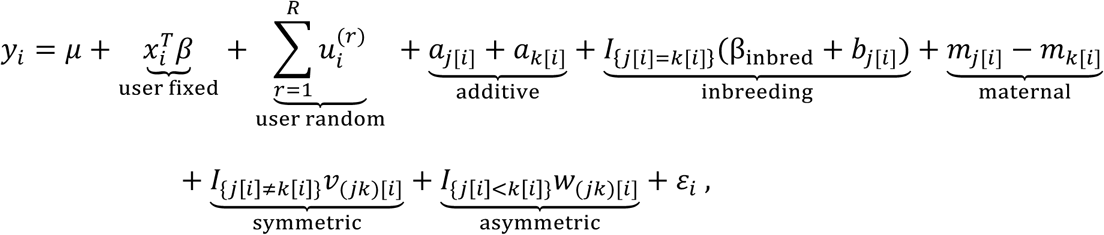

where *j*[*i*], *k*[*i*] and (*jk*)[*i*] respectively denote the female, male and female-male combination relevant to individual *i*, each group of effects parameters is modeled from its own random effects distribution, e.g. *a*_*j*_~*N*(0, *τ*_*a*_^2^) fixed covariates *x_i_*, and *R* additional random-effect components are included as *u_i_*^(*r*)^~*N*(0, *τ*_*a*_^2^), ∀ *r* ∈ { 1, …, *R*}. Planting density (1-3 scale, 1=low, 3=high) was included as a fixed covariate to capture linear trends and as a random effect to estimate deviations from linearity. Year (CA2015, WI2015, CA2016) was also included as a random effect. This model was fitted in BayesDiallel using a MCMC Gibbs sampler with 5 chains, 10000 iterations, and a burn-in of 1000.

Although convention is to model heterosis as a dominance effect, this method instead models inbred-specific deviations from heterozygote-based predictions; that is, homozygotes (i.e. parental selfs), which are a minority in the diallel, are treated as a special class. Thus, additive effects are a dosage effect of parent *j* in combination with another sampled parent. Epistasis, modeled as deviations from additivity, is represented by symmetric and asymmetric effects. Symmetric effects model differences specific to a given cross, regardless of parental inheritance and independent of reciprocal effects (i.e. crosses *jk* and *kj* have the same effect). Asymmetric effects model deviations from symmetric effects due to differences between reciprocal crosses (i.e. *jk* and *kj* have different effects).

### Diallel variance projection as a heritability-like measure

In order to report the relative contribution of each diallel inheritance class to a given phenotype, Crowley et al. (2014) proposed the diallel variance projection (VarP). This approach uses the posterior predictive distribution of effects from BayesDiallel to simulate future, complete, perfectly balanced diallels of the same parental lines. In each simulated dataset, the contribution of each inheritance class (additive, inbreeding, etc.) is then calculated as its sum of squares (SS) divided by the total phenotype SS. The resulting proportion, the VarP, is similar to the traditional heritability of Mather and Jinks (1982) and Lynch and Walsh (1998) but with two important differences: 1) it is explicitly prospective, in that it seeks to describe how much additive effects, say, would impact a future experiment; and 2) its estimation is more precise, since it is calculated as a function primarily of the effects parameters (e.g. *a*_1_,*a*_2_*, …,a*_6_), which are well informed by the data, rather than of the variance components (*τ*_*a*_^2^, *τ*_*m*_^2^, etc.), which are typically not. VarPs are calculated from multiple posterior draws leading to a complete posterior distribution of the VarP for each inheritance class, summarized here as highest posterior credibility intervals. Credibility intervals that include zero are interpreted as not contributing positive, nonzero information to the prediction of *y_i_*, whereas credibility intervals excluding zero provide strong evidence that an effect is important to the model.

## RESULTS

### Imputation of missing data

There was a high incidence of missing data due to variation in seed production and disease pressure (Table 1, **Figure S3**). A large proportion of missing data occurred in the WI2015 environment, which was subject to severe infestation by Alternaria leaf blight, a fungal pathogen that causes leaf necrosis and plant death in carrots (Pryor and Strandberg 2001). Distributions of imputed data matched those expected from observed data when accounting for environmental variation (**Figures S5-6**).

### Additive and non-additive gene action contributed to observed phenotypes

Most phenotypes were positively correlated and significant at *α*=0.001, with the exception of the ratio for shoot:root biomass with both canopy height and width at 80 DAP (Figure 2). Griffing’s analysis revealed significant genotypic differences for all phenotypes (Table 2), which are reflected in posterior predicted means from BayesDiallel (Figure 3 a, b; **Figures S7-12**). For all traits, both GCA and SCA contributed significantly to the observed genotypic variation, suggesting that both additive and non-additive effects are important. The ratio of GCA to SCA varied for each trait but was less than one for all phenotypes except shoot:root ratio, indicating a more prominent role of non-additive gene action for most traits (Table 2). This is reaffirmed by the results from BayesDiallel, in which the highest posterior density intervals for inbreeding effects were greater and more dispersed than for additive effects (Figure 3 c, d; **Figures S13-19**).

**Figure 2.**
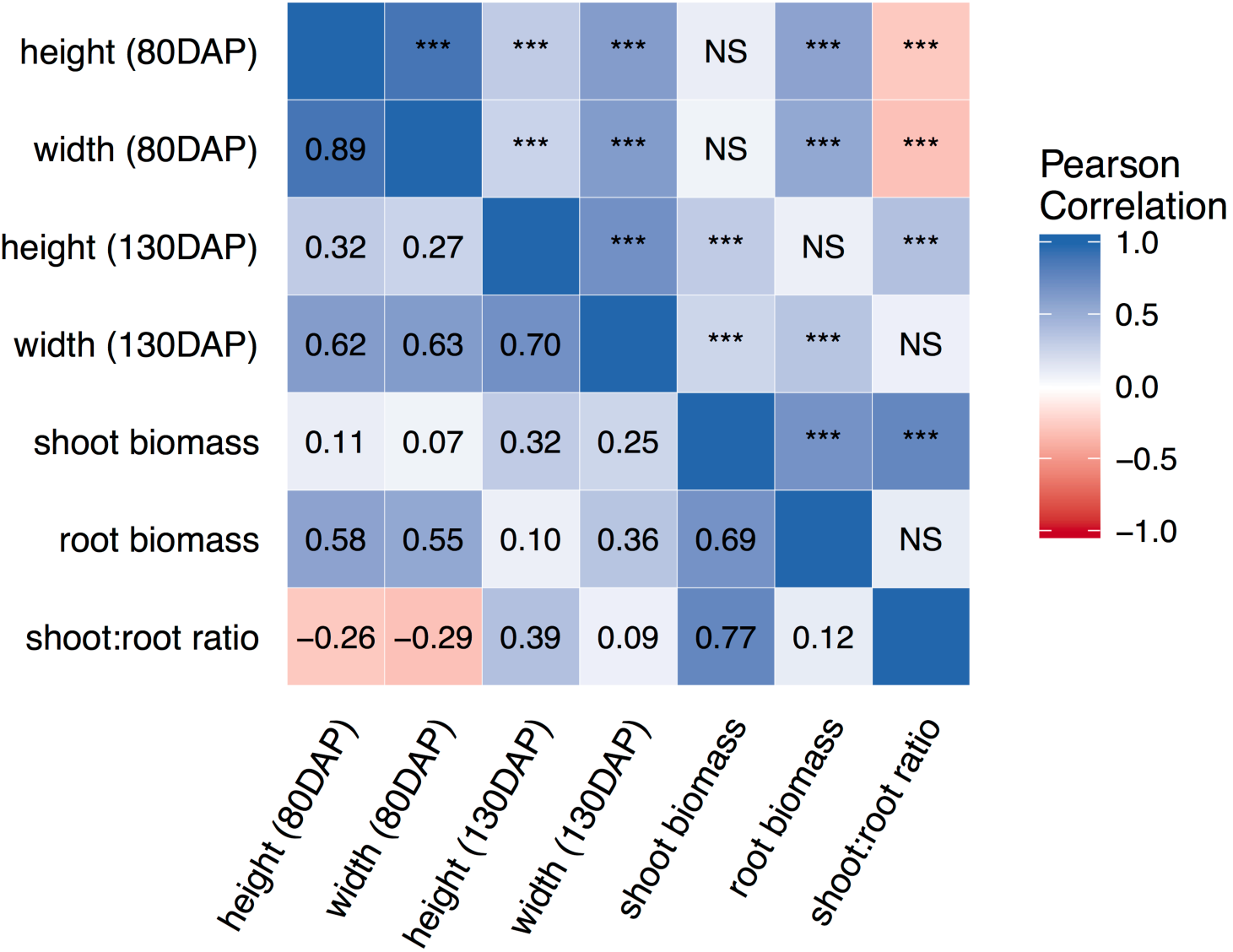
Pearson’s correlations (lower diagonal) and significance (upper diagonal) among carrot growth traits measured in this study. Significance codes: *** *P* ≤ 0.001, ** *P* ≤ 0.01, * *P* ≤ 0.05, NS not significant.

**Figure 3.**
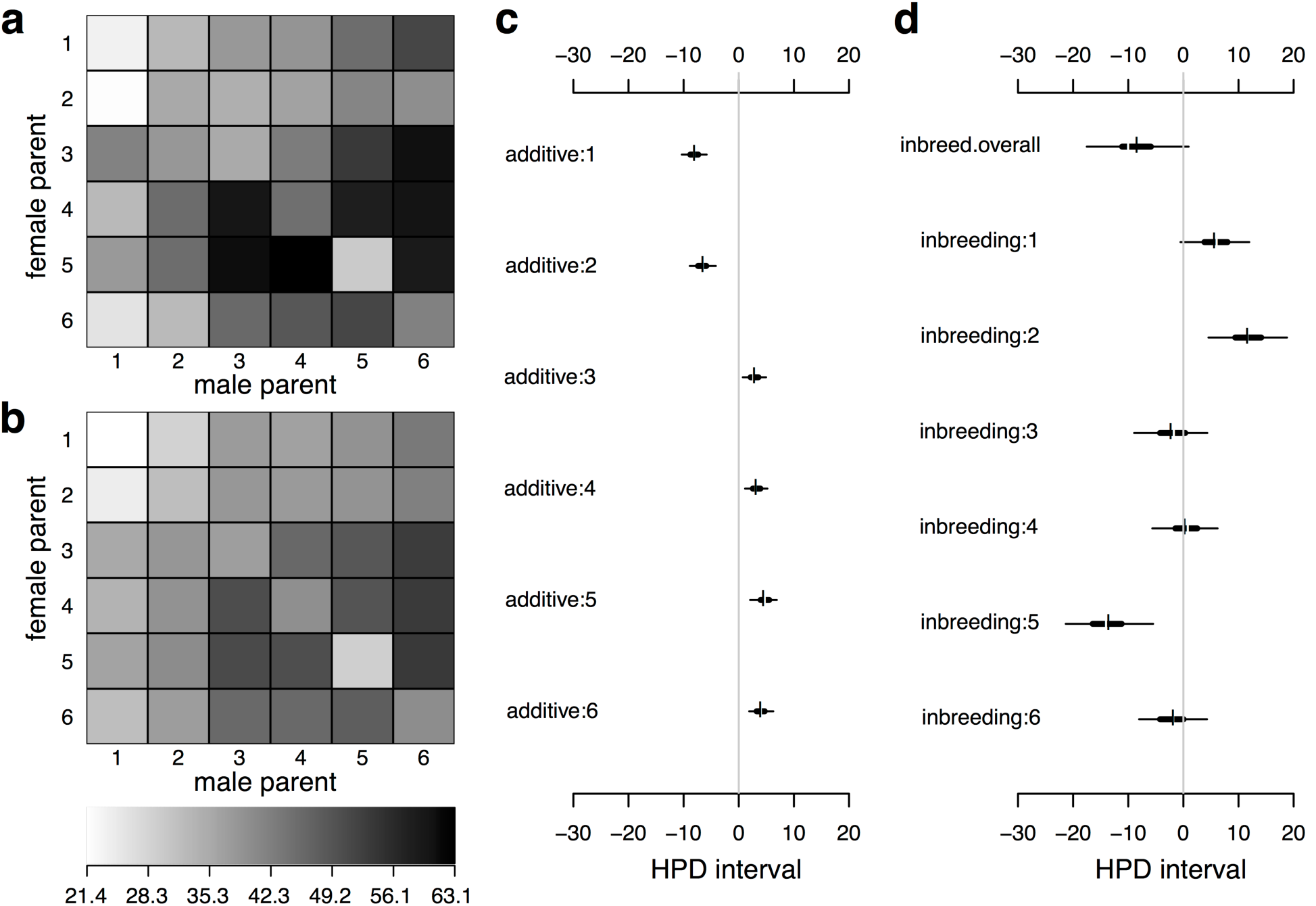
Diallel effects for carrot canopy height (cm) at 130 days after planting. **a** and **b** show observed and predicted means, respectively. Shading indicates height on a scale from 21.4 cm (lighter) to 63.1 cm (darker). Posterior predicted means in **b** are the result of fitting the model in BayesDiallel to the data in **a**. The highest posterior density (HPD) intervals for additive effects are shown in **c** and the HPD intervals for inbreeding effects are shown in **d**. For each effect, thin and thick horizontal lines show the 95% and 50% HPD intervals, respectively, and the vertical break displays the posterior mean. Key: 1 = L6038, 2 = L7550, 3 = P0159, 4 = Nbh2189, 5 = P6139, 6 = B7262.

**Table 2.**
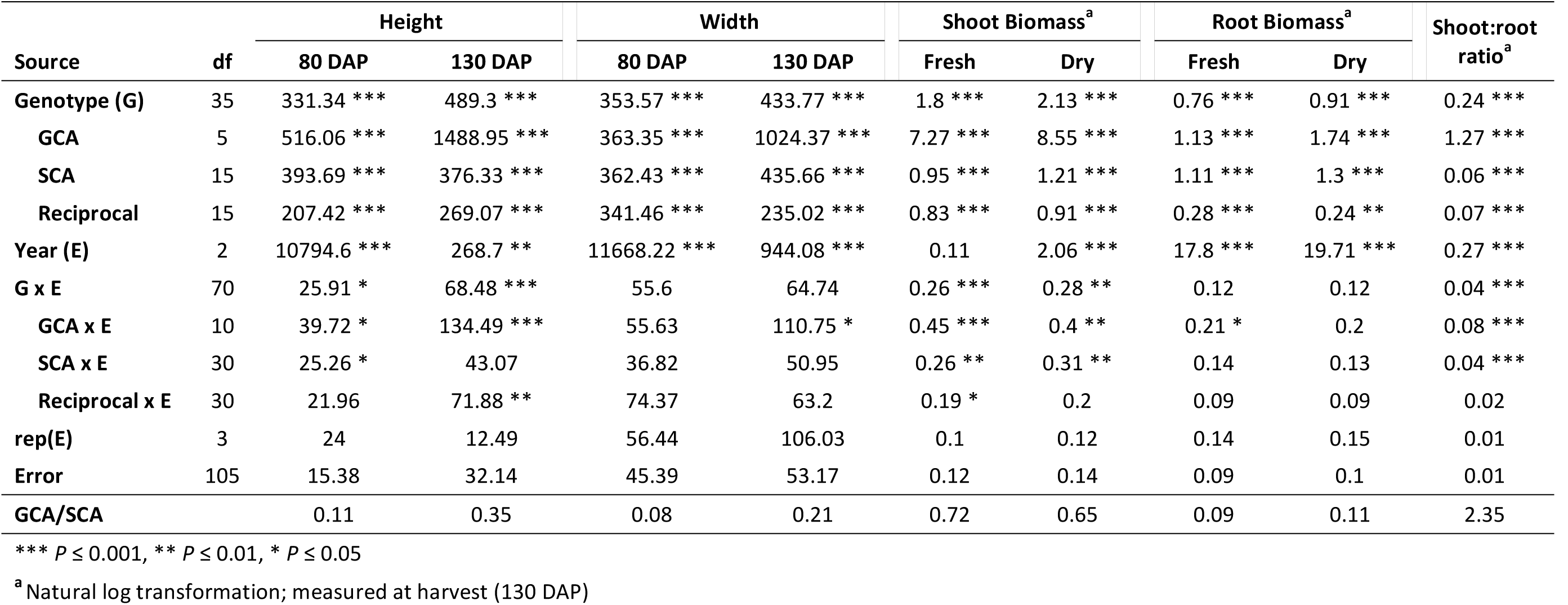
Griffing’s Method I, Model I ANOVA mean square values for carrot growth traits, including: canopy height and width (cm), shoot biomass (g), root biomass (g), and the ratio of shoot:root biomass. The ratio of general combining ability (GCA) to specific combining ability (SCA) variance is also reported.

### Inbreeding effects differed across genetic backgrounds

Results from Griffing’s analysis indicated that the observed phenotypes, with the exception of shoot:root ratio, were largely under non-additive genetic control (Table 2), which is also reflected in the posterior predicted means and highest posterior density intervals from BayesDiallel (**Figures S7-19**). These effects are illustrated by canopy height at 130 DAP, for which inbred lines were an average of 8.4 cm shorter than their hybrid counterparts (overall inbreeding, *β_inbred_*, in Figure 3). Additionally, the intensity of inbreeding also varied across genetic background (Figure 3, **Figures S13-19**). Relative to heterozygotes, line L6038 had a minimal net reduction in canopy height of 3.0 cm (*β_inbred_ + b_j_*_[1]_), while line B7262 had a net 22 cm reduction in canopy height (*β_inbred_ + b_j_*_[5]_) (Figure 3).

### Identification of superior parents, hybrids, and testers for applied breeding

GCA estimates were compared to determine the relative performance of each parent (Table 3). Parent L6038 had negative and significant GCA for all traits except canopy height and width at 80 DAP. Low and significant GCA was also observed in parent L7550 for height (130 DAP), shoot biomass, and the ratio of shoot:root biomass. For canopy height, parents with positive and significant GCA included Nbh2189 (130 DAP), P6139 (80 and 130 DAP), and B7262 (130 DAP). Parent Nbh2189 was the only inbred with significant and positive GCA for canopy width (130 DAP). Parents P0159 and B7262 had high and significant GCA for both shoot biomass and the ratio of shoot:root biomass. Positive and significant GCA for root biomass was only observed for parent P0159.

**Table 3.** Pooled estimates of general combining ability (GCA) for carrot growth traits combined across all growing environments.

| Parent | Height |  | Width |  | Biomass<sup>a</sup> |  |  |
| --- | --- | --- | --- | --- | --- | --- | --- |
|  | 80 DAP | 130 DAP | 80 DAP | 130 DAP | Shoot | Root | Shoot:root ratio |
| **L6038** | -1.39 | -6.5*** | 0.09 | -3.5** | -0.53*** | -0.19* | -0.19*** |
| **L7550** | -2.83 | -4.94*** | -0.63 | -2.23 | -0.15 | -0.03 | -0.08** |
| **P0159** | 0.85 | 1.43 | -1.56 | -1.41 | 0.39*** | 0.27*** | 0.1*** |
| **Nbh2189** | 2.43 | 3.62** | 3.15 | 7.13*** | 0.11 | 0.06 | 0.03 |
| **P6139** | 3.52* | 3.87** | 1.86 | 0.53 | -0.15 | -0.07 | -0.03 |
| **B7262** | -2.58 | 2.52* | -2.92 | -0.53 | 0.33*** | -0.03 | 0.18*** |
| **Grand Mean** | 32.27 | 45.77 | 43.57 | 53.74 | 1.55 | 2.55 | 0.6 |

\*\*\* $P \leq 0.001$ , \*\* $P \leq 0.01$ , \* $P \leq 0.05$
<sup>a</sup> Natural log transformation; dry weight as measured at harvest (130 DAP)

GCA estimates largely agree with the results from BayesDiallel, which provided similar rankings based on posterior predicted means (Figure 3 a, b; **Figures S7-12**) and HPD intervals (Fig. 3 c, d; **Figures S13-19**). For canopy height (130 DAP), hybrids with parents L6038 and L7750 were, on average, about 6.5 cm shorter, while hybrids with parents P6139 and B7262 were an average of 5.0 cm taller. (Figure 3). The observed and posterior predicted means for canopy height (130 DAP) also demonstrate relatively higher values for hybrids with parents P6139 and B7262, as well as lower values for hybrids containing parents L6038 and L7550 (Figure 3 a, b).

SCA effects were identified as crosses that performed better or worse than expected based on the GCA values of the contributing parents (Table 4). Hybrid Nbh2189 x P6139 had high SCA for both height and width (80 and 130 DAP). For shoot biomass, the largest SCA was observed in hybrid Nbh2189 x B7262. Hybrids with high SCA for root biomass included L7550 x B7262, P0159 x Nbh2189, and Nbh2189 x B7262. No significant positive effects were observed for shoot:root ratio. Line Nbh2189 was the most discriminating parent in hybrid combination for all phenotypes except root biomass, suggesting it can serve as a valuable tester for shoot growth in applied breeding.

**Table 4.**
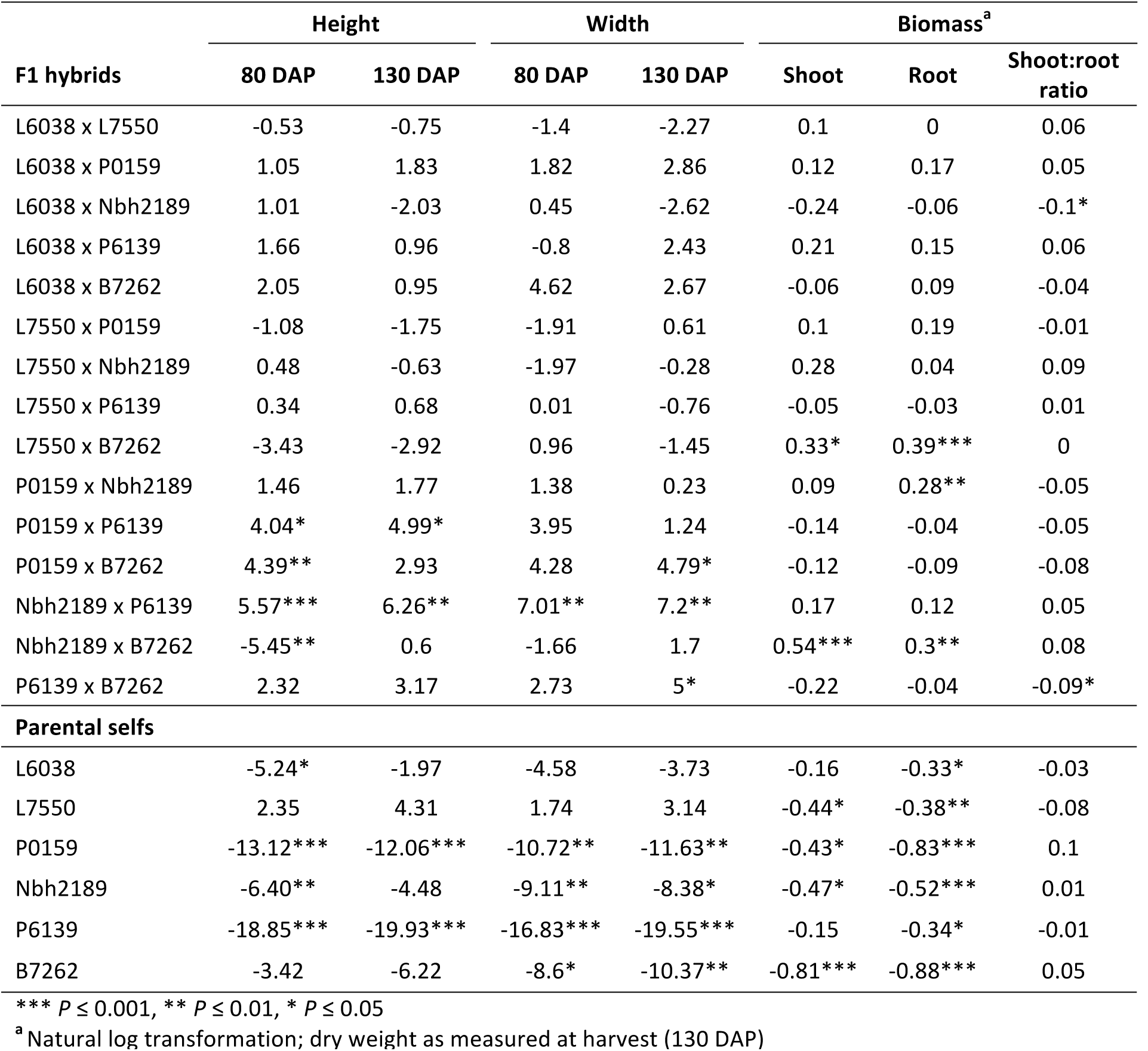
Pooled estimates of specific combining ability (SCA) for carrot growth traits combined across all growing environments.

**Table 5.**
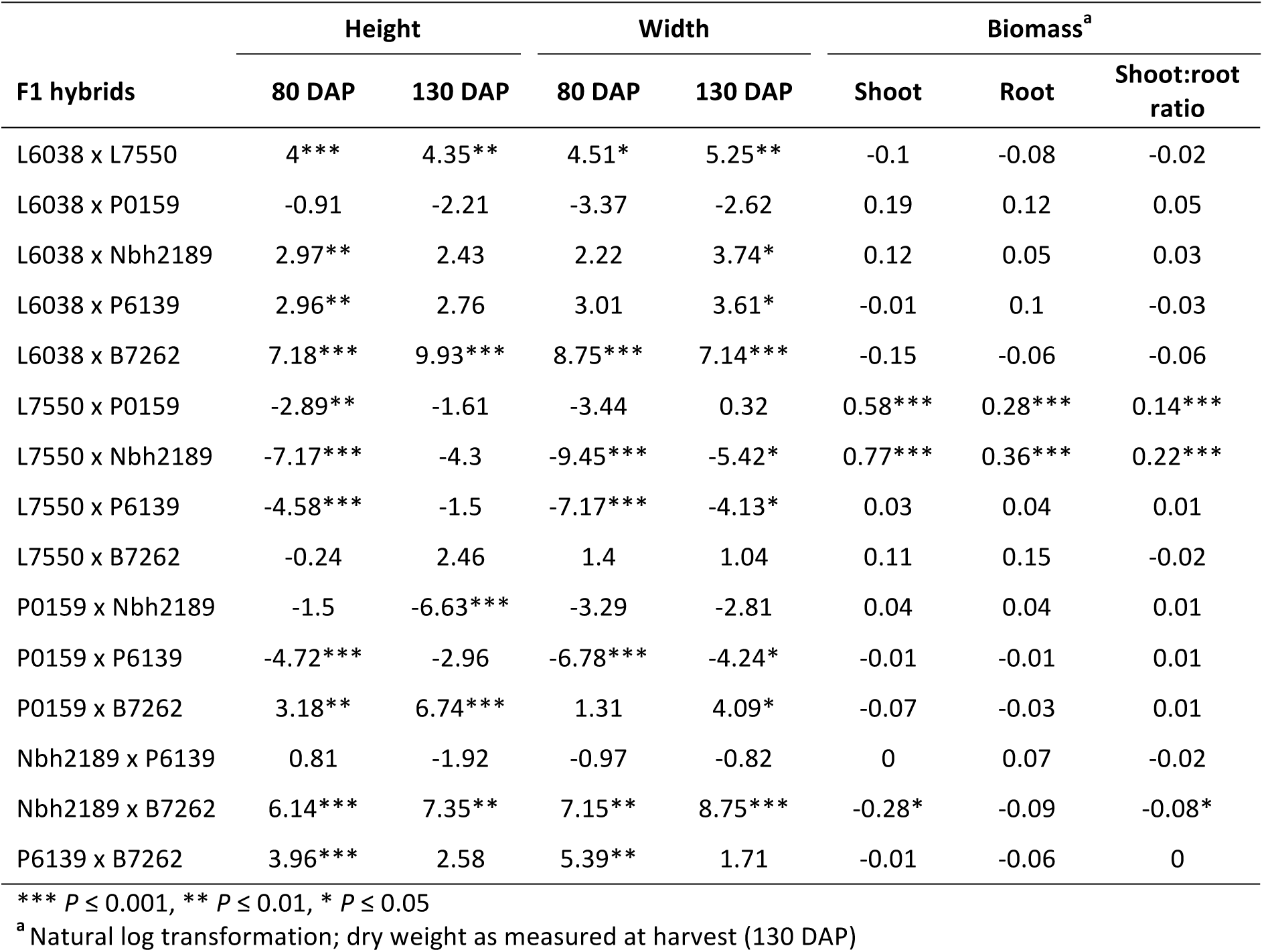
Pooled estimates of reciprocal cross effects for carrot growth traits over all growing environments.

| F1 hybrids | Height |  | Width |  | Biomass<sup>a</sup> |  |  |
| --- | --- | --- | --- | --- | --- | --- | --- |
|  | 80 DAP | 130 DAP | 80 DAP | 130 DAP | Shoot | Root | Shoot:root ratio |
| L6038 x L7550 | 4*** | 4.35** | 4.51* | 5.25** | -0.1 | -0.08 | -0.02 |
| L6038 x P0159 | -0.91 | -2.21 | -3.37 | -2.62 | 0.19 | 0.12 | 0.05 |
| L6038 x Nbh2189 | 2.97** | 2.43 | 2.22 | 3.74* | 0.12 | 0.05 | 0.03 |
| L6038 x P6139 | 2.96** | 2.76 | 3.01 | 3.61* | -0.01 | 0.1 | -0.03 |
| L6038 x B7262 | 7.18*** | 9.93*** | 8.75*** | 7.14*** | -0.15 | -0.06 | -0.06 |
| L7550 x P0159 | -2.89** | -1.61 | -3.44 | 0.32 | 0.58*** | 0.28*** | 0.14*** |
| L7550 x Nbh2189 | -7.17*** | -4.3 | -9.45*** | -5.42* | 0.77*** | 0.36*** | 0.22*** |
| L7550 x P6139 | -4.58*** | -1.5 | -7.17*** | -4.13* | 0.03 | 0.04 | 0.01 |
| L7550 x B7262 | -0.24 | 2.46 | 1.4 | 1.04 | 0.11 | 0.15 | -0.02 |
| P0159 x Nbh2189 | -1.5 | -6.63*** | -3.29 | -2.81 | 0.04 | 0.04 | 0.01 |
| P0159 x P6139 | -4.72*** | -2.96 | -6.78*** | -4.24* | -0.01 | -0.01 | 0.01 |
| P0159 x B7262 | 3.18** | 6.74*** | 1.31 | 4.09* | -0.07 | -0.03 | 0.01 |
| Nbh2189 x P6139 | 0.81 | -1.92 | -0.97 | -0.82 | 0 | 0.07 | -0.02 |
| Nbh2189 x B7262 | 6.14*** | 7.35** | 7.15** | 8.75*** | -0.28* | -0.09 | -0.08* |
| P6139 x B7262 | 3.96*** | 2.58 | 5.39** | 1.71 | -0.01 | -0.06 | 0 |

\*\*\* $P \leq 0.001$ , \*\* $P \leq 0.01$ , \* $P \leq 0.05$
<sup>a</sup> Natural log transformation; dry weight as measured at harvest (130 DAP)

### Non-genetic factors and interactions influenced observed variation

Highly significant reciprocal effects were detected for all traits in Griffing’s analysis (Table 2), suggesting parent-of-origin influences phenotypic expression. For increasing height and width (80 and 130 DAP), lines L6038 and P0159 tended to perform best as female parents and lines L7550 and B7262 tended to perform best as male parents. Significant increases were also observed for shoot biomass, root biomass, and shoot:root ratio when line L7550 was used as a female parent and when lines P0159 and Nbh2189 were used as male parents.

Genotype by environment interaction (GxE) was significant for canopy height (80 and 130 DAP), shoot biomass (fresh and dry), and shoot:root ratio (Table 2). For corresponding traits with significant GCAxE, SCAxE, and ReciprocalxE interactions, estimates for each year are provided in **Tables S2-4**. Significant GCAxE interactions were observed for canopy height (80 and 130 DAP), shoot biomass, and shoot:root ratio. For canopy height (130 DAP), GCA ranked consistently negative across environments for parents L6038 and L7550 (Figure 4). Parent Nbh2189 had positive GCA in all environments, but effects were only significant for the CA2015 and CA2016 locations (**Table S2**). In general, relative rankings for parents remained consistent across environments, but the ability to distinguish significant effects was diminished in the WI2015 environment. The performance of parents P0159 and B7262 was notably inconsistent and fluctuated between negative and positive values of GCA (Figure 4). SCAxE interactions were significant for height (80 DAP), shoot biomass, and shoot:root ratio, but it was still possible to identify consistently high performing hybrids across environments (**Table S3**). Similarly, significant ReciprocalxE interactions were observed for canopy height (130 DAP) and fresh shoot biomass (**Table S4**). Differences across replications within a year were not significant.

**Figure 4.**
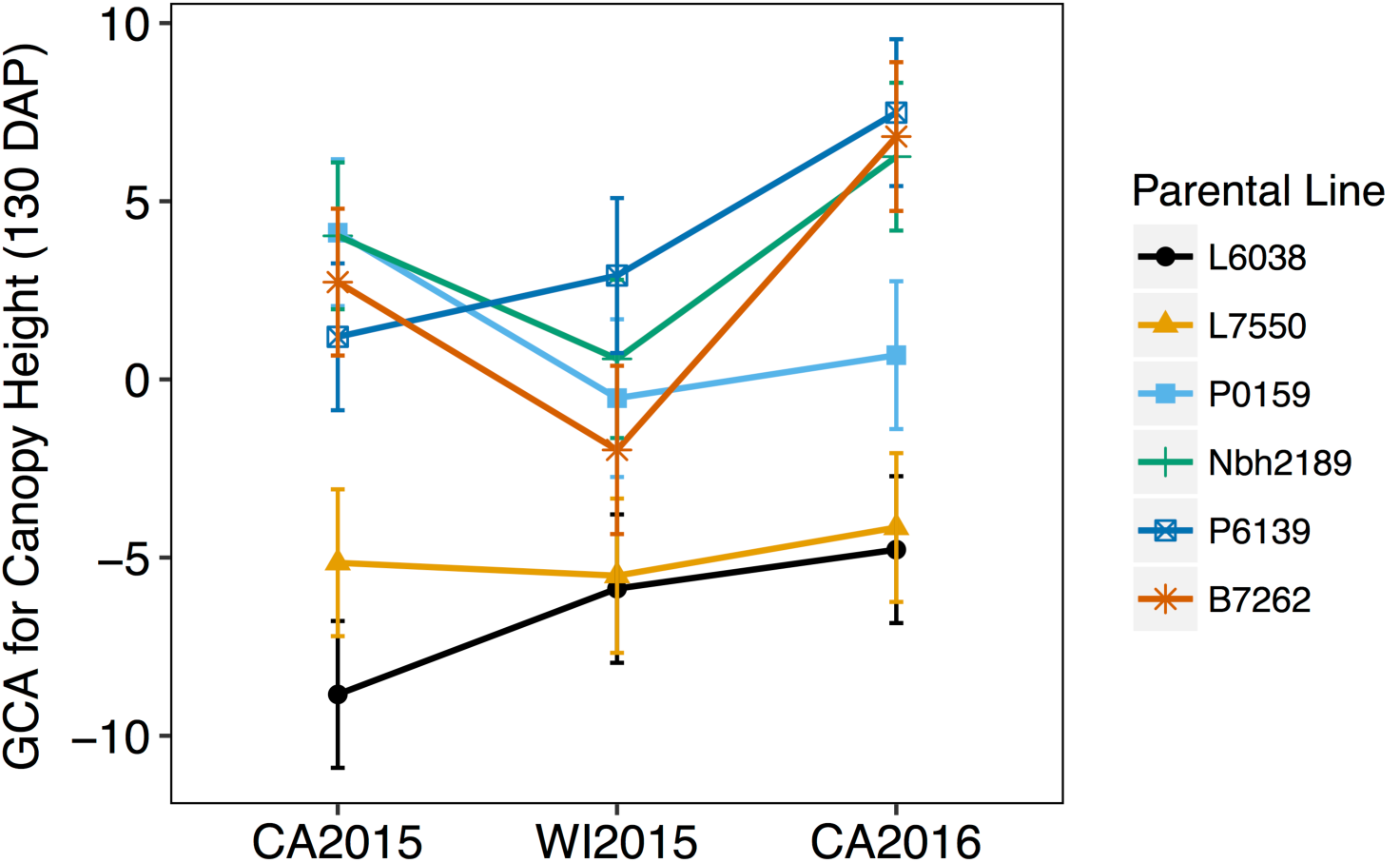
Interaction of general combining ability (GCA) and year (WI2015, CA2015, CA2016) for canopy height at 130 DAP.

Additional effects estimated in BayesDiallel included planting density (fixed and random) and year (random) (**Figure S20**). On average, planting density increased plant height in a mostly linear fashion, with a greater effect at 80 DAP (5.39 cm) compared to 130 DAP (3.51 cm) (Figure 5). Similarly, year had a greater influence at 80 DAP than at 130 DAP, with the highest mean in the WI2015 season and the lowest mean in the CA2016 season (Figure 5).

**Figure 5.**
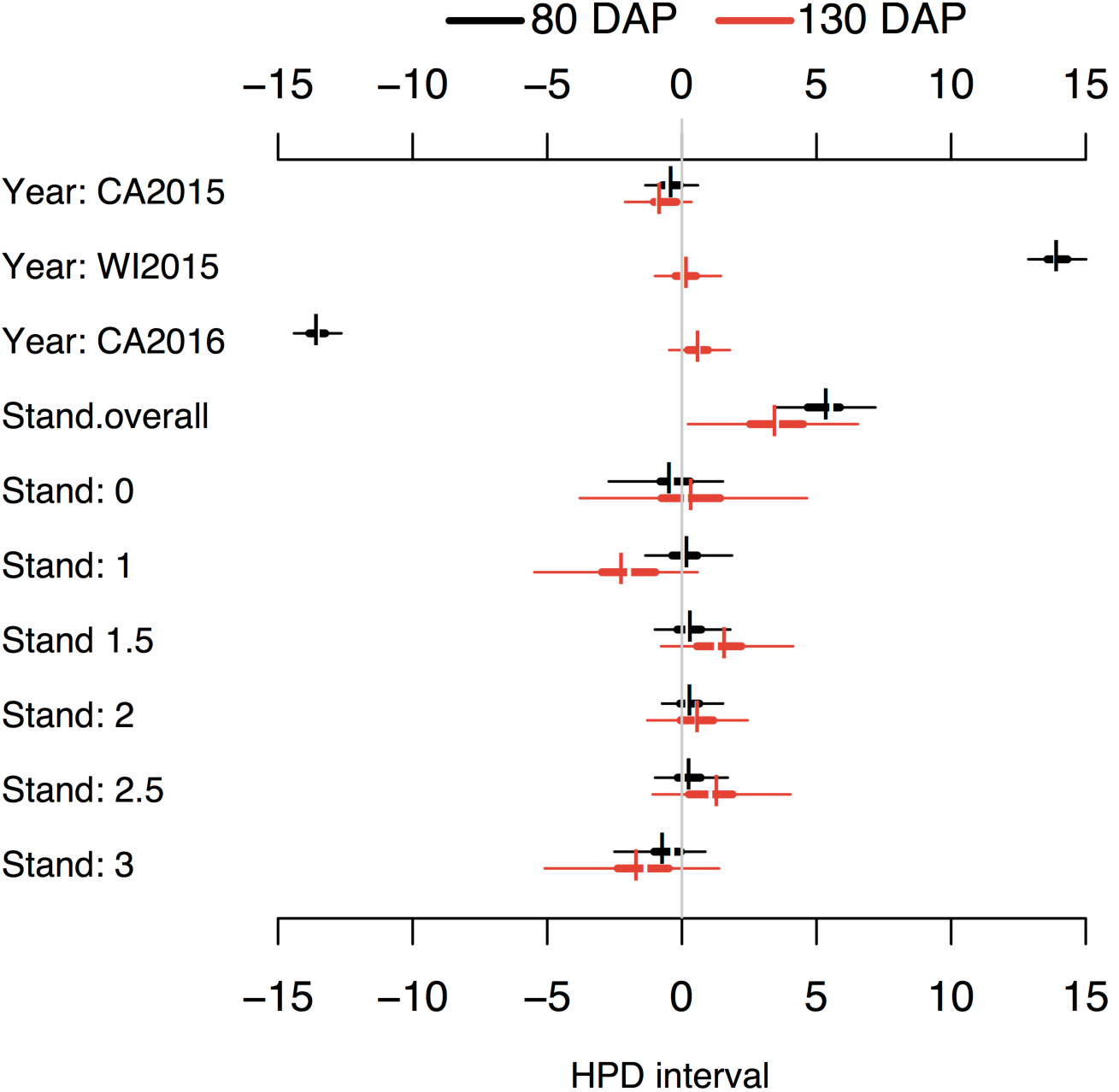
Highest posterior density (HPD) intervals of year and planting density (stand; 1-3 scale, 1 = low, 3 = high) for canopy height at 80 DAP (black) and 130 DAP (red). For each effect, thin and thick horizontal lines show the 95% and 50% HPD intervals, respectively, and the vertical break displays the posterior mean.

### Genetic architecture varied across traits and over developmental time

Although most correlations among phenotypes ranged from moderate (r>0.3) to high (r>0.5) (Figure 2), genetic architecture varied substantially by trait and across developmental time (Figure 6). For canopy height and width, the GCA/SCA ratio increased approximately three-fold from 80 to 130 DAP, demonstrating a higher proportion of non-additive variation early in the growing season (Table 2). Results from BayesDiallel exhibit a similar relationship, with overall inbreeding (*β_inbred_*) explaining more variation than additive effects for height and width at 80 DAP (Table 6, Figure 6).

**Figure 6.**
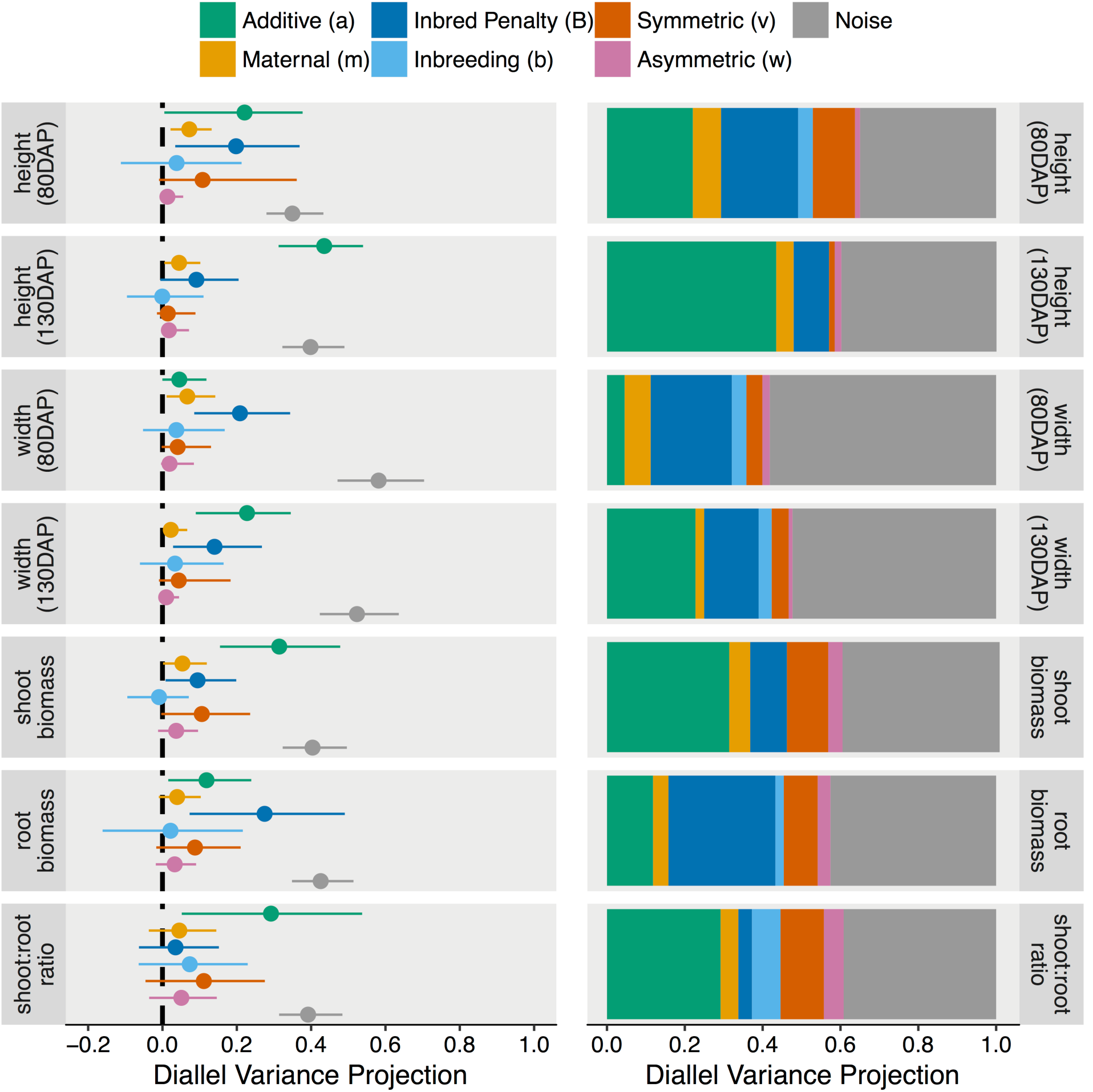
Diallel variance projections characterizing the genetic architecture for each trait, including additive (a), maternal (m), overall inbreeding penalty (B), parent-specific inbreeding (b), symmetric (v), and asymmetric (w) effect classes. **Left:** mean and 95% credibility intervals of effect classes for each trait. **Right:** Mean values showing overall genetic architecture.

**Table 6.**
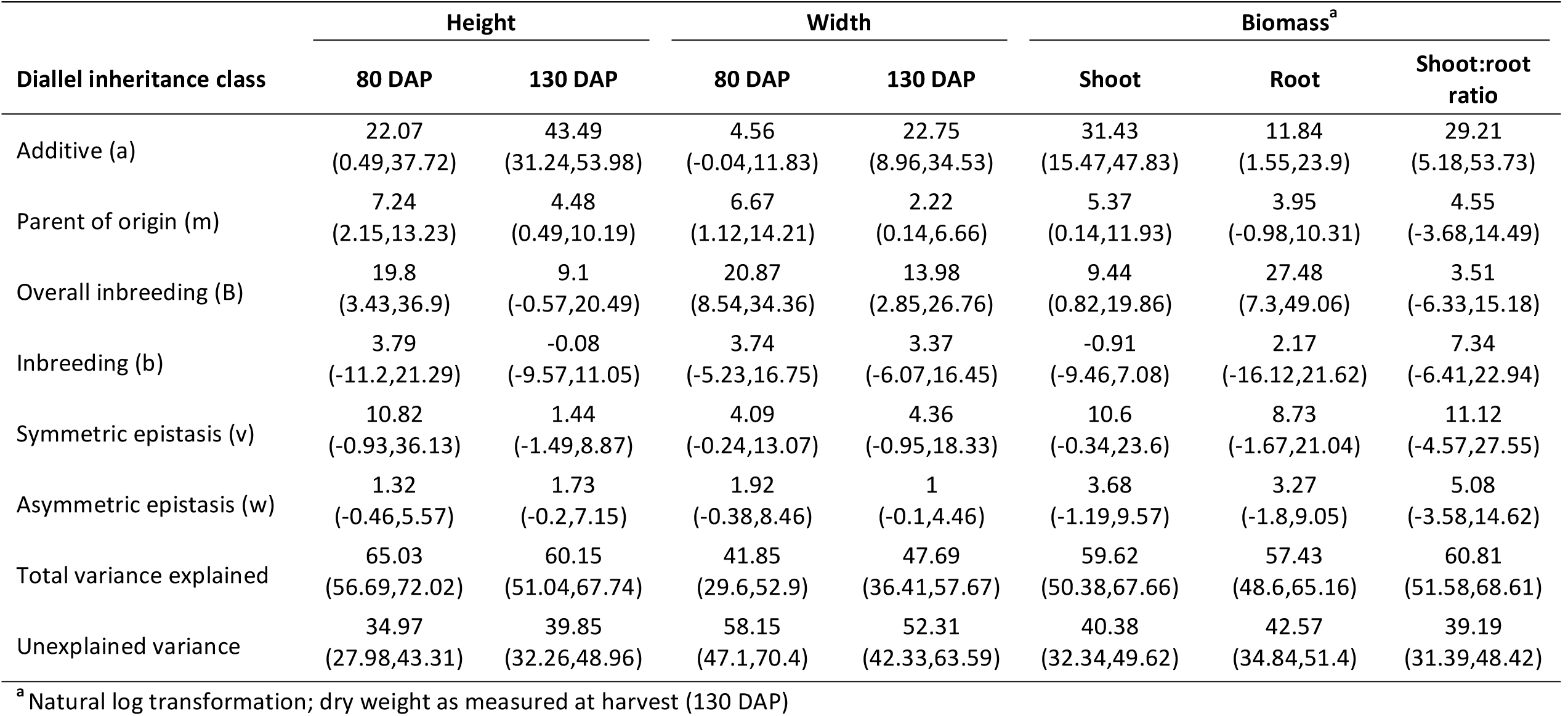
Diallel variance projection (VarP) for carrot growth traits. Under each trait (column) is listed the predicted percentage and 95% credibility intervals of variance that would be attributable to each class of effect. Predictions are conditional on a future complete diallel composed of the same parental lines.

As described byCrowley et al. (2014), the variance projection of the additive diallel inheritance class, VarP[a], can be likened to narrow sense heritability (*h^2^*). Traits with significant additive effects included canopy height at 80 DAP (VarP[a] = 0.22), canopy height at 130 DAP (0.43), canopy width at 130 DAP (0.23), shoot biomass (0.31), root biomass (0.12), and shoot:root ratio (0.29) (Table 6, Figure 6).

The influence of non-additive variation was largely due to overall inbreeding, which contributed significantly to canopy height at 80 DAP (VarP[B] = 0.20), canopy width at 80 DAP (0.21) and at 130 DAP (0.14), shoot biomass (0.09), and root biomass (0.27) (Table 6, Figure 6). However, parent-specific inbreeding effects, symmetric epistasis, and asymmetric epistasis did not contribute significantly to the predicted phenotypes (Table 6, Figure 6). While parent-of-origin is not a genetic effect, it did explain variation for canopy height at 80 DAP (Var[m] = 0.07) and 130 DAP (0.04), canopy width at 80 DAP (0.07) and 130 DAP (0.02), and shoot biomass (0.05) (Table 6, Figure 6).

## DISCUSSION

### Primary gene action

In this study, we estimated genetic, parent-of-origin, and environmental effects on carrot growth traits for six carrot inbred lines and their combinations in a 6x6 diallel framework. Significant genetic variation contributed to all carrot growth attributes, suggesting that there is potential to improve these traits in carrot.

Apart from canopy width at 80 DAP, all phenotypes had some level of predictability based on inbred performance, as evidenced by the presence of non-zero additive effects. Traits with high additivity included canopy height (130 DAP) and shoot biomass, both of which are well documented as highly heritable polygenic traits that play a fundamental role in plant fitness and adaptation (Khush 2001; Meyer et al. 2004; Peiffer et al. 2014).

For the parental lines in this study, we observed varying sensitivity to inbreeding, which could be due to genetic divergence and/or differing levels of prior inbreeding (East 1936; Birchler et al. 2003). This matches expectations based on the biological constraints of outcrossing in carrot, which has putative susceptibility to inbreeding depression (Simon 2000). Consequently, hybrid vigor was evident for canopy height (80 DAP), canopy width (80 DAP), and root biomass, which had high proportions of non-additive genetic variation and significant estimates of inbreeding, suggesting that causative loci may have heterozygote advantage (Charlesworth and Willis 2009). This result coincides with widespread evidence of heterosis in plants, whereby hybrids demonstrate increased developmental speed and greater biomass relative to their inbred parents (East 1936; Birchler et al. 2003; Meyer et al. 2004), and agrees with previous observations of hybrid vigor for root weight in carrot (Simon et al. 1982).

### Breeding strategies

With discovery of cytoplasmic male sterility (CMS) in carrot (Welch and Grimball 1947), breeding strategies transitioned from selection in open-pollinated populations to an inbred-hybrid system, thereby improving crop uniformity and vigor (Peterson and Simon 1986). We expect that traits with a significant contribution of inbreeding, such as canopy height (80 DAP), canopy width (80 DAP), and root biomass, will be responsive to commonly used hybrid breeding strategies in carrot, such as reciprocal recurrent selection. Alternatively, selection for traits with high additivity, such as canopy height (130 DAP), canopy width (130 DAP), and shoot biomass, may allow more rapid genetic gain while indirectly selecting for positively correlated traits under non-additive control. For all phenotypes in this study, we identified promising parental lines and hybrids for use in applied breeding. Additionally, line Nbh2189 was notable as a favorable tester for future breeding efforts on shoot growth.

Accounting for genotype by environment interaction (GxE) is especially important in biennial crops like carrot, as breeding programs rely on winter nurseries to achieve an annual breeding cycle (Simon and Goldman 2007). We conducted trials in CA and WI, which are two of the leading carrot production regions and representative of common, but contrasting, breeding environments. In general, significant GxE interactions did not affect the ability to identify high and low rankings among parents and hybrids. Thus, we anticipate that environmental differences are important, but should have a minimal impact on selection efforts.

### Source-sink relationships

Biomass partitioning between the shoot and root is a major consideration in carrot breeding and has been extensively studied, both in a breeding and an ecological context (Barnes 1979; Currah and Thomas 1979; Hole et al. 1983; Thomas et al. 1983; Hole and Dearman 1991). The ideotype for carrot shoot growth is rapid initial growth that plateaus following canopy closure, simultaneously reducing the critical weed free period and promoting taproot development. Equally important is avoiding growth habits with large, dense canopies, which foster a microclimate that is conducive to the development of foliar diseases (Simon et al. 2008).

Consistent with findings by Hole et al (1983), we found a strong linear relationship between the log transforms of shoot and root biomass (r = 0.69, *P*<0.001). However, the ratio of shoot:root biomass had a wide range across parents (0.19-0.72), providing evidence that high shoot biomass is not necessary to produce roots with high biomass, and vice versa.

Previous work has demonstrated rapid and early acquisition of dry matter in carrot storage roots, with the taproot constituting 42% of the plant dry weight at 67 days after sowing (Benjamin and Wren 1978). Interestingly, our results demonstrate that canopy height and width at 80 DAP were negatively correlated with the ratio of shoot:root biomass (r = −0.26 and −0.29, respectively, *P<* 0.001) and positively correlated with root biomass at harvest (r = 0.58 and 0.55, respectively, *P*<0.001). Conversely, canopy height at harvest (130 DAP) was positively correlated with shoot:root ratio (r = 0.39, *P*<0.001) and not significantly correlated with root biomass (r = 0.10, *P*=0.18). This suggests that early shoot growth is important for root biomass accumulation and agrees with previous conclusions that these traits are subject to hybrid vigor.

### Method of analysis

When applied appropriately, traditional diallel analysis provides valuable information on combining abilities for parental lines and the underlying gene action for complex traits (Griffing 1956a). However, the benefits of diallel mating designs are often overshadowed by practical and theoretical constraints of the analysis. The requirement for complete, balanced data is especially restrictive, particularly in crops with poor seed set and limited availability of inbred lines. In this study, we performed classical evaluation using Griffing’s Method I and a modern analysis using a general Bayesian approach (BayesDiallel). Both methods provided similar conclusions regarding primary gene action, parental rankings, and hybrid performance.

Although the underlying model is more complex, BayesDiallel provided numerous practical advantages over Griffing’s analysis, which are thoroughly discussed by Lenarcic et al (2012). Of relevance in this experiment was the robustness of BayesDiallel to missing data, which was a substantial challenge when applying Griffing’s model and is pervasive in field experiments. For the latter, we chose to address missing data using multiple imputation, which produces a set of plausible values to replace missing information (Rubin 1987). However, there are several caveats and compromises regarding multiple imputation in that there are inadequate or vague diagnostics and, although simple in principle, methods to pool multi-factor ANOVA results are often vague or are not widely accessible (van Ginkel and Kroonenberg 2014; Grund et al. 2016). Furthermore, the ability of BayesDiallel to analyze data sets in the presence of missing data allows breeders to leverage data from partial diallels, which are often generated by proxy in a breeding program.

A notable advantage of BayesDiallel was the option to add covariates for year and planting density as random and fixed effects. Posterior distributions for year matched expectations based on observed data, with higher means observed in WI2015 compared to CA2015 and CA2016. The inclusion of planting density was especially advantageous and matched expectations from previous studies, which demonstrated significant effects of planting density on canopy height and biomass partitioning in carrot (Bleasdale 1967; Benjamin and Sutherland 1992; Traka-Mavrona 1996; Li et al. 1996)

Precise estimates of heritability are useful when determining which breeding strategy will result in the most genetic gain. While it is possible to estimate heritability through variance component decomposition using Griffing’s model II, this method requires that the parents are selected from a random mating population (at least according to its traditional interpretation; see discussion in Lenarcic et al. 2012), and is not accompanied by a measure of estimation uncertainty. As an alternative, the diallel variance projection (VarP) in BayesDiallel has benefit for applied breeding by (1) describing how several inheritance classes will influence future experiments composed of the same parents, and (2) providing a 95% credibility interval as a measure of uncertainty, which affords more flexibility when designing future experiments and estimating potential for genetic gain.

### Conclusions

The rise of sustainable agriculture has tasked breeders to develop cultivars with improved weed competitive ability. Using traditional and modern approaches, we analyzed diallel data to describe the quantitative inheritance of previously uncharacterized traits in carrot, which have been demonstrated to confer improved competitive ability in other crops. However, future trialing for weed competitive ability in carrot will be essential to validate the utility of these traits, to determine the underlying mechanism of competitive ability (i.e. tolerance or suppression), and to assess relative fitness costs (e.g. trade-offs with yield).

Results from this study support applied breeding efforts for carrot shoot growth in numerous ways, most notably through the quantification of inbred performance, the identification of a useful tester line (Nbh2189), and the assessment of potential hybrid combinations. Furthermore, the detailed characterization of the inbred parents in this study provides a foundation for the development of a multi-parental advanced generation intercross (MAGIC) population in carrot, which will facilitate future in-depth genetic studies (Huang et al. 2015).

Lastly, we demonstrate the utility of the BayesDiallel framework for modeling heritable and non-heritable components of carrot phenotypes. This example will make BayesDiallel more accessible as a resource in the plant breeding community to maximize the potential of diallel experiments, especially in under-resourced crops.

## Acknowledgements

We are grateful to Dr. Shelby Ellison, Guillaume Ramstein, Joe Gage, and Robert Corty for helpful discussion regarding the content of this manuscript and to Rob Kane for technical support. This research was supported by the USDA National Institute of Food and Agriculture under award number 2011-51300-30903 of the Organic Agriculture and Research Extension Initiative.

## LIST OF SUPPLEMENTARY FIGURES

**Figure S1**. Carrot inbred lines used as parents in this study.

**Figure S2.** Parental means and 95% confidence intervals for all traits.

**Figure S3**. Incidence of missing data by replication and year for canopy height (130 DAP).

**Figure S4**. Convergence of imputed Markov chain Monte Carlo (MCMC) chains for all traits.

**Figure S5**. Distributions of observed and imputed values for all traits.

**Figure S6**. Distributions of observed and imputed values for selected traits by year.

**Figures S7-12**. Observed and predicted means for each trait evaluated.

**Figures S13-19**. Highest posterior density (HPD) intervals of inheritance classes for each trait evaluated.

**Figure S20**. Highest posterior density (HPD) intervals of random effect classes for all traits evaluated.

## LIST OF SUPPLEMENTARY TABLES

**Table S1.** Pooled results of Griffing's ANOVA (Method I, Model I) for multiply imputed data of carrot growth traits.

**Table S2.** Estimates of general combining ability (GCA) by year (CA2015, WI2015, CA2016) for carrot growth traits with significant genotype by environment interactions (GxE) and significant GCAxE interactions.

**Table S3.** Estimates of specific combining ability (SCA) by year (CA2015, WI2015, CA2016) for carrot growth traits with significant genotype by environment interactions (GxE) and significant SCAxE interactions.

**Table S4.** Estimates of reciprocal effects by year (CA2015, WI2015, CA2016) for carrot growth traits with significant genotype by environment interactions (GxE) and significant ReciprocalxE interactions.

